# Zebrafish glial-vascular interactions progressively expand over the course of brain development

**DOI:** 10.1101/2024.09.27.615488

**Authors:** Lewis G. Gall, Courtney M. Stains, Moises Freitas-Andrade, Bill Z. Jia, Nishi Patel, Sean G. Megason, Baptiste Lacoste, Natasha M. O’Brown

## Abstract

Glial-vascular interactions are critical for the formation and maintenance of brain blood vessels and the blood-brain barrier (BBB) in mammals, but their role in zebrafish is not well understood. Our previous work has detailed the timeline of BBB functional maturation in zebrafish, revealing a conserved mechanism of BBB induction through the suppression of endothelial transcytosis. Yet, as opposed to extensive research on glial-vascular interactions in rodents, such interactions remain largely overlooked in the zebrafish model system. Here, we focus on glial-vascular development in the zebrafish brain, leveraging three glial gene promoters: *gfap* (glial fibrillary acidic protein), *glast* (an astrocyte-specific glutamate transporter), and *glastini* (a new, shortened, equally effective version of the Glast promoter). Using these glial promoters, sparse labeling revealed fewer glial-vascular interactions during early larval stages, with both glial coverage and contact area increasing as the zebrafish brain matured. We then generated stable transgenic lines for both the Glast and Glastini promoters and observed similar increases in glial coverage during larval development, starting at ∼30% coverage at 3 days post-fertilization (dpf) and peaking at ∼60% at 10 dpf. Ultrastructural assessment of glial-vascular interactions using electron microscopy (EM) confirmed a progressive increase in glial coverage over larval development, with maximal coverage reaching ∼70% in adult zebrafish, significantly lower than the nearly 100% coverage observed in mammals. Finally, immunogold-EM labeling confirmed that cells identified as glia in aforementioned morphological analyses were indeed Glast-positive. Taken together, our results identify the temporal profile of glial-vascular maturation in the zebrafish brain.

## Introduction

Proper neural function requires a precisely regulated microenvironment carefully maintained by a specialized layer of vascular endothelial cells that form the blood-brain barrier (BBB). Past studies have shown the importance of distinct signaling pathways between endothelial cells and surrounding cell types, such as pericytes, astrocytes, neurons, and microglia, in inducing and maintaining barrier properties^1–7^. Vascular support cells, or mural cells, called pericytes share a basement membrane with these endothelial cells and are required for both the induction of BBB properties and maintenance of these properties in adulthood^5,6,8^. Astrocytes, which are born mostly postnatally in mice, after the initial embryonic establishment of the BBB, completely ensheath the vasculature with polarized endfeet in the brain and have been shown to be required for maintenance of BBB properties^4,7,9^.

Zebrafish have recently emerged as a powerful model system to examine the dynamics of the BBB in vivo, with many evolutionarily conserved molecular and cellular regulators of BBB function^2,10–13^. However, while it has been established that zebrafish possess astrocyte-like cells in the spinal cord with similar morphological, spatial and dynamic calcium signaling to mammalian astrocytes^14^, whether or not these glial cells play the same role in the zebrafish BBB remains unknown. Zebrafish glia express many of the same hallmark genes to mammalian astrocytes, including *aldh1l1, aqp4, gfap, glast (slc1a3), glt1 (slc1a2)*, and *s100ß* (*s100b*)^14–16^. However, unlike mammalian AQP4, which is concentrated at astrocytic endfeet surrounding blood vessels^17,18^, AQP4 in adult zebrafish spans the entire length of the glial process^16^, similar to its distribution in immature postnatal mammalian astrocytes^18^. Zebrafish astroglia appear to contact the vasculature as early as 7 dpf^15^. However, it remains unclear how these glial-vascular interactions develop and whether glial coverage reaches comparable levels to those seen in the mature mammalian brain. Thus, further investigation is needed to fully understand the dynamics of glial-vascular coverage in the developing zebrafish brain.

In this context, we sought to characterize glial-vascular interactions developmentally first through mosaic transgenic labeling of glia with two different promoters, *gfap* and *glast*, to assess possible differences between the two labeled glial populations, revealing low levels of glial coverage early in development. We then extended these analyses to juvenile and adult stages, revealing further increases in glial-vascular contacts, and instances glial ensheathment of the vasculature in the adult, as observed throughout the mammalian brain. We also generated a condensed version of the original *glast* promoter, titled *glastini*, which we found drives indistinguishable expression in glia. Furthermore, we generated and analyzed two new stable lines, *glast:myrGFP* and *glastini:myrGFP*, which revealed developmental increases in total glial-vascular coverage over larval development. Finally, using electron microscopy (EM), we confirmed an increase in glial endfoot coverage of the vasculature with age, with maximal coverage in adults, correlating well with our fluorescent analyses. Our results show a maximum glial coverage of 70% on the vasculature in the adult zebrafish brain, compared to nearly 100% in mice, thus highlighting species-specific differences in glial-vascular maturation.

## Results

### Mosaic glial analyses reveal increased vascular contacts with age

To assess glia in zebrafish, we first generated mosaic glial transgenics in a Tg(*kdrl:mCherry*) background using the previously established *gfap:EGFP* plasmid^21^, which labels both immature and differentiated glial cells. We then allowed these F0 mosaics to develop to different larval stages and examined their glial-vascular interactions beginning at 3 dpf, before the BBB is functional^10^, until 10 dpf, well after the BBB is fully functional (Fig. 1A). These analyses revealed strong GFP labeling throughout the brain, and an abundance of different morphological subtypes of GFP+ cells, including rounded amoeboid cells (Fig. 1B), those with long radial processes (Fig. 1C), as well as several branched, astroglial-like cells (Fig. 1D). We also noticed that several of these cells directly interacted with the vasculature.

**Fig. 1.**
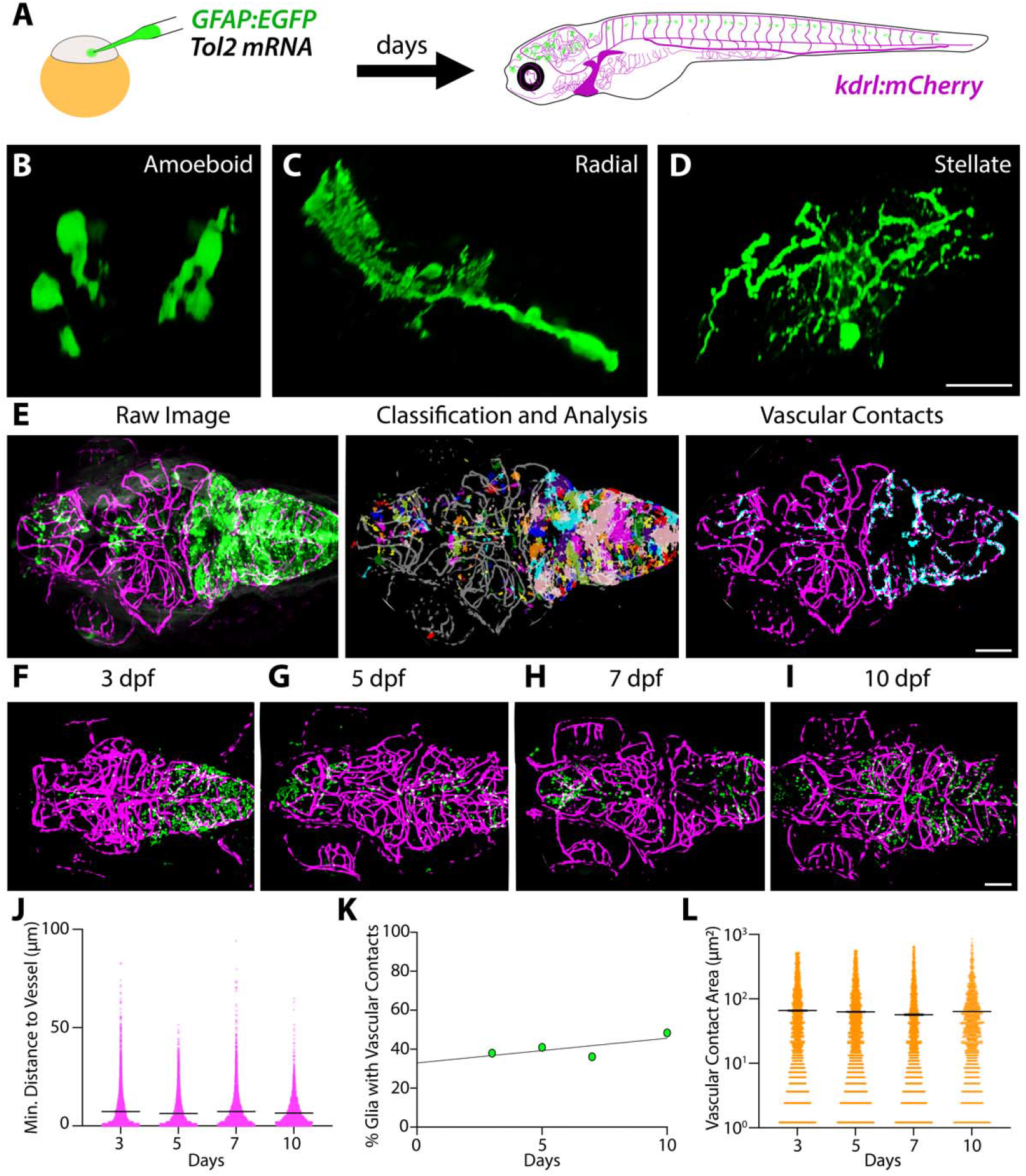
Analysis of gfap+ glial-vascular interactions over larval development. (A) Schematic of experimental design to generate mosaically labeled glia using the GFAP:GFP plasmid and tol-2 transgenesis. (B-D) 3D projections of individual glia, revealing 3 different subtypes of glial cells that were labeled by the gfap promoter: amoeboid (B), radial (C), and stellate (D). (E) Example image and resulting classification and analysis leading to the vascular contacts mask through our custom pipeline. (F-I) Representative maximum intensity projections of the mosaically labeled glia (green) and vasculature (magenta) at 3 (F), 5 (G), 7 (H), and 10 dpf (I). (J-L) Quantification of the mean distance to the nearest blood vessel for each glial cluster (J), the % of total measured glia with any vascular contacts (K), and the glial-vascular contact area (L) reveal relatively stable glial-vascular interactions over time. Scale bars represent 10 μm (D) and 50 μm (E, I).

To quantify these features, we employed a custom Python pipeline to extract glial morphological data and assess glial-vascular contacts over the course of zebrafish development (Fig. 1E-I). These analyses revealed a relatively consistently low minimum distance to the nearest blood vessel of less than 10 μm away (Fig. 1J), with about 45% of *gfap*+ glia having vascular contacts throughout larval development (Fig. 1K). The size (area) of these vascular contacts also remained relatively steady with an average of 62 μm^2^ (Fig. 1L).

To further investigate these interactions, we next assessed terminally differentiated *glast*+ glia using the same analysis pipeline. Unlike *gfap*+ glia, *glast*+ glia displayed temporally regulated vascular interactions, with decreased distances to the nearest blood vessels and increased vascular contacts with age and a slight increase in contact area from 7 to 10 dpf that stayed consistent at 14 dpf (Fig. 2). Given this increase throughout larval development, we extended these *glast* analyses through to adulthood, and observed much larger increases in glial-vascular contacts; however, we never observed more than 70% of glia having any vascular contact (Fig. 2J). Our analyses of cleared adult brains showed a single instance of glial ensheathment of the vasculature in the forebrain (Fig. 2H), indicating that zebrafish glia have the capacity for vascular wrapping, as typically seen by astrocytes in adult mice.

**Fig. 2.**
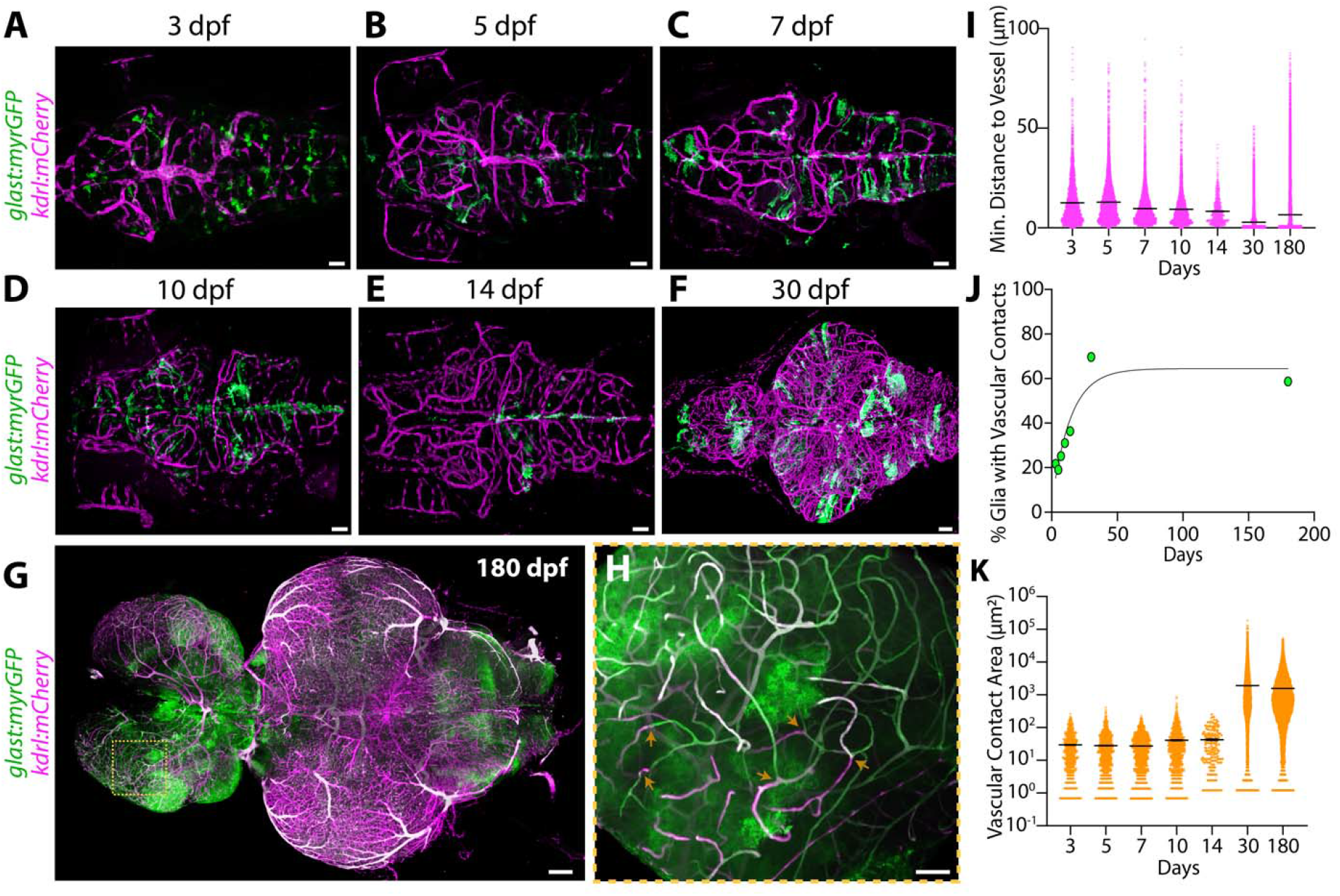
Glast+ glial-vascular interactions increase with age. (A-F) Representative 3D projections of glast+ glial (green) and vascular (magenta) masks at 3 (A), 5 (B), 7 (C), 10 (D), 14 (E), and 30 dpf (F). (G-H) Maximum intensity projection (MIP) of an entire cleared adult zebrafish brain revealed glial ensheathment of the vasculature in 1 out of 14 adult fish in the boxed area of the forebrain (H). There are multiple breakpoints (orange arrows) of glial coverage, revealing discrete domains of vascular coverage. (I-K) Quantification of the mean distance to the nearest blood vessel for each glial cluster (I), the % of total measured glia with any vascular contacts (J), and the glial-vascular contact area (K) reveal an increase in the amount of glia with vascular contacts and the size of these contacts with age. Scale bars represent 50 μm (A-E, H), 100 μm (F), and 200 μm (G).

### New, shorter ‘*glastini*’ promoter maintains glial expression

While working with the original 9.55 kb *glast* promoter (Fig. 3A), we noticed that it had lower efficiency in labeling glia compared to the *gfap* promoter. Considering this may be partly due to the larger size of the promoter, we aimed to remove extraneous sequences while maintaining glial expression. This led to the creation of a 3 kb promoter we call ‘*glastini*’, that takes 2 non-adjacent sequences from the original *glast* promoter with putative open chromatin, including the 5’ UTR for *slc1a3b*, and concatenates them (Fig. 3A). Mosaic injections with the *glastini* promoter revealed radial and stellate glial labeling (Fig. 3B), similar to what we had previously observed with the original *glast* mosaic injections^14^. Upon running our analysis pipeline and comparing *glast*+ glia to *glastini*+ glia at 5 dpf, we did not observe any significant difference in the glial contact areas between the two promoters (Fig. 3). This indicates that, despite dramatically shortening the promoter sequence, glial expression was not compromised.

**Fig. 3.**
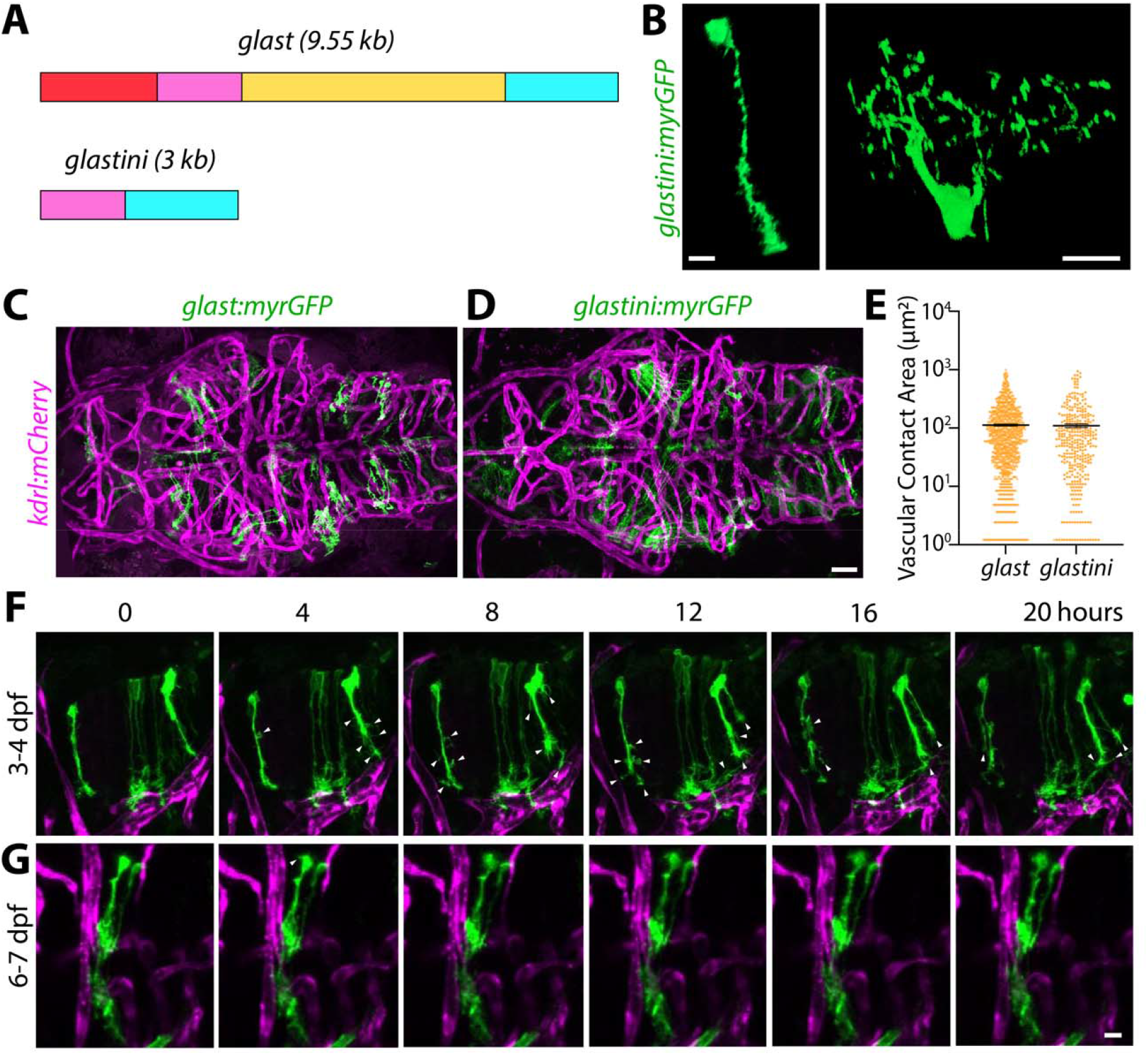
*Glastini*, a condensed *glast* promoter, retains glial expression and highlights developmental glial dynamics. (A) Schematic of the original *glast* promoter and the condensed *glastini* promoter with the conserved areas color-coded accordingly. The turquoise sequence contains the entire 5’ UTR and the pink sequence contains a downstream H3K27ac peak. (B) 3D image of different individual glial subtypes labelled by the *glastini* promoter, radial and stellate. (C) MIP of a mosaic *glast* injected 5 dpf fish showing glial cells labeled with *glast* (green) interacting with brain vasculature (magenta). (D) MIP of a mosaic *glastini* injected 5 dpf fish, showing similar glial cells labeled with *glastini* (green) interacting with brain *kdrl+* vasculature (magenta). (E) Python analysis of the two different glial promoters, *glast* and *glastini*, reveals a similar vascular contact area for glial cells labeled by both promoters (p=0.78 by unpaired t-test). (F-G) Still images from time-lapse imaging of *glastini+* glia at 3 dpf (F, Video S1) reveals highly dynamic glial protrusions (white arrowheads) in regions without vascular contacts and more stable glial-vascular interactions at 6 dpf (G, Video S2). Scale bars represent 5 μm (B, G) and 50 μm (D).

Using this *glastini* promoter, we performed long-term time-lapse imaging to assess dynamic changes in glial-vascular interactions. At 3 dpf, we observed many glial processes extending towards and contacting the vasculature, as well as sending projections into the surrounding area (Fig. 3F, Video S1). Furthermore, projections making vascular contacts were maintained longer than those that did not. When repeating imaging at 6 dpf, we observed an overall reduction in projection number and motility, and stable vascular contacts (Fig. 3G, Video S2).

### Analysis of total glial coverage reveals developmental increases

Next, we generated stable transgenic lines for *glast* and *glastini*. Both lines drove similar expression throughout the central nervous system (CNS; Fig. 4). Using these lines, we performed a comprehensive analysis of total glial coverage of the vasculature during larval development. Our analyses of the *glast:myrGFP* line revealed minimal glial coverage (30%) at 3 dpf, with no significant change from 3 to 5 dpf. However, by 7 dpf, there was a substantial increase to about 50%, which remained high at 10 dpf (58%). Similarly, the *glastini:myrGFP* line showed low coverage at 3 dpf (33%), followed by a larger increase at 5 dpf (46%), and similarly high levels at 7 dpf (53%) and 10 dpf (60%). The corresponding data between both lines further suggest that the new *glastini* promoter drives glial expression that is indistinguishable from the original *glast* promoter. While our analyses were done in the whole brain, we noted that the forebrain and midbrain had higher glial coverage than the hindbrain, with regions near the midline remaining completely devoid of glial contacts, even at 10 dpf.

**Fig. 4.**
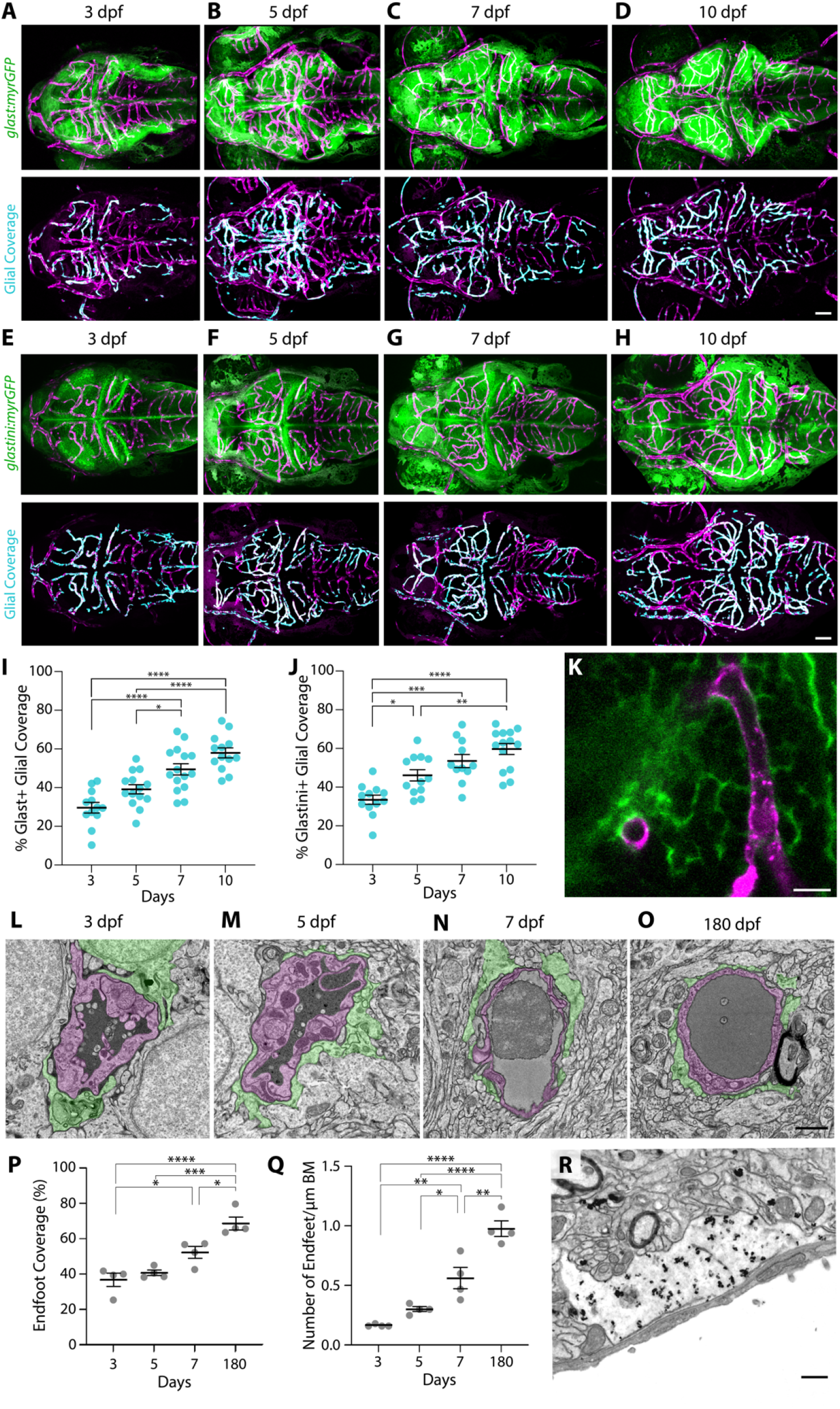
Stable glial transgenic lines reveal a steady increase in glial coverage of the vasculature with age. (A-D) Representative MIP images of stable *glast:myrGFP* glial– vascular interactions from 3 (A), 5 (B), 7 (C), and 10 dpf fish (D). Raw images are on top with glast:myrGFP+ glia (green) and kdrl:mCherry vasculature (magenta). The vascular contact mask (turquoise) is imposed on the vascular signal in the lower panels. (E-H) Representative MIP images of stable *glastini:myrGFP* glial–vascular interactions from 3 (E), 5 (F), 7 (G), and 10 dpf fish (H). Raw images are on top with glastini:myrGFP+ glia (green) and kdrl:mCherry vasculature (magenta). The vascular contact mask (turquoise) is imposed on the vascular signal in the lower panels. (I-J) Quantification of the percentage of glial coverage of the vasculature reveals a steady increase from 3dpf to 10dpf in both *glast:myrGFP* (I) and *glastini:myrGFP* (J) transgenic lines. (K) Zoomed in 2 μm thick MIP image reveals multiple points of direct glial contact of the vasculature and multiple vascular regions completely devoid of glial coverage in a 7 dpf fish. (L-O) Zoomed out TEM images with the vessels pseudocolored magenta and the glial endfeet pseudocolored green in 3dpf (L), 5 dpf (M), 7 dpf (N), and 180 dpf adult brains (O). (P-Q) Quantification of glial coverage of the brain vasculature (P) and number of endfeet per μm of the vascular basement membrane (Q) reveals a steady increase of glial coverage with age. Each dot represents the average value of 10 capillaries analyzed for a single fish (N=4). (R) Immunogold for glast:GFP+ glia in adult brains confirms glast+ glial endfoot contacts with the vasculature, as marked by the dark, silver-enhanced gold particles within the electron light endfoot. Scale bars represent 50 μm (D,H), 10 μm (K), 1 μm (O) and 500 nm (R).. ****p<0.0001, ***p<0.001, **p<0.01, *p<0.05 by one-way ANOVA with multiple comparisons.

Finally, as a complementary approach, we examined glial-vascular interactions at the ultrastructural level using transmission electron microscopy (TEM) during larval development (3-7 dpf) and in adults (Fig. 4L-O). TEM analyses revealed similar larval glial coverage levels to our fluorescent analyses in the stable lines (Fig. 4I & J), with 37% coverage at 3 dpf, 41% coverage at 5 dpf, and 52% coverage at 7 dpf (Fig. 4P). Glial coverage was highest in the adults at 69%. When we looked at the number of endfeet per μm of endothelial basement membrane, we observed a steady increase in the number of endfeet contacting the vasculature with age (Fig. 4Q), further suggesting that glial-vascular interactions enhance with brain maturation. Finally, we confirmed that the cells morphologically annotated as glial cells in TEM images were in fact *glast*+, using immunogold labelling for GFP in adult *glast:myrGFP* fish. These analyses revealed strong, highly-selective expression of the GFP reporter in electron-light endfeet surrounding endothelial cells, as well as in other glial processes throughout the neuropil (Fig. 4R).

## Discussion

In this study, we rigorously characterized the development of glial-vascular interactions in zebrafish using a variety of approaches, including mosaic and stable transgenic glial lines and electron microscopy, all of which revealed consistent patterns of increased glial coverage of the vasculature with age. To ensure unbiased analysis, we developed a custom Python pipeline for 3D image analysis that identifies individual glial features from our mosaic injections, such as morphology, surface area, volume, and the number and size of glial contacts with the vasculature. This pipeline could be applied to analyze various cell-cell interactions where at least one cell population is sparsely labeled, such as glial-neuronal or neuronal-vascular interactions. It may also help explore and quantify more complex cell-cell interactions, both in normal development and in disease or mutant states. Additionally, we generated *glastini*, a truncated *glast* promoter that retains the same glial expression patterns as *glast*, even in a stable transgenic line. Since *glastini* is one-third the size of *glast*, this innovation allows more space for larger genes to be expressed specifically in glia, providing increased flexibility for transgenic expression in zebrafish glia.

The zebrafish model system is gaining momentum for studying CNS blood vessels and neurovascular interactions. Although zebrafish share a molecularly and cellularly conserved vascular unit with endothelial cells and pericytes^2,10,12,13,27^, the nature and extent of glial-vascular interactions in zebrafish remained poorly characterized and required further investigation. Our analyses revealed that at most, in the adult, zebrafish have about 70% glial coverage of the vasculature. This is in stark contrast to mice, which have 99.8% coverage by postnatal day 14 (P14)^25,28^, indicating that there are differences in the interactions between zebrafish glial cells and the brain vascular versus in mammals, even in the most mature state. In addition to these changes in maximal glial coverage of the vasculature, zebrafish glia also have differential localization of the water channel Aquaporin-4 (Aqp4) compared to mice. While adult mouse astrocytes localize AQP4 expression to their endfeet surrounding the vasculature, zebrafish glia express Aqp4 throughout the entire cell body^16,29^. Interestingly, in mammals, astrocytes initially express AQP4 throughout the entire cell body during early development, but this expression becomes restricted to the endfeet as glial coverage reaches 100% by P14^18^. This might suggest that zebrafish glia maintain a more “immature” molecular phenotype that could be resolved with some cross-species transcriptomic comparisons over the course of glial maturation in both zebrafish and mice.

AQP4 has been previously studied as a key player in waste clearance^30^ and its localization has been linked to improved clearance within the mammalian brain^31^. This raises the possibility that waste clearance pathways in adult zebrafish may be less efficient or use different mechanisms than in the adult mouse brain, though further research in zebrafish is needed to confirm this. Notably, the adult zebrafish brain is significantly smaller in volume (64x) compared to the adult mouse brain^32^. It is possible that a larger brain requires more advanced waste clearance mechanisms, whereas in a smaller brain, passive mechanisms might suffice. Comparative studies of glial-vascular interactions and waste clearance in mammals with varying brain sizes could offer deeper insights into the relationship between brain size and the need for glial-vascular coverage in efficient waste clearance.

With this in mind, one might question whether incomplete or immature glial coverage of the brain vasculature offers any advantages. Zebrafish are renowned for their ability to regenerate many tissues including the heart, fins, brain, and spinal cord^33^. We believe there may be a link between the extent of glial-vascular interactions and neural tissue regeneration. For example, neonatal mice (P2-P4) can recover from cryogenic brain injury, but this regenerative ability is completely lost by P14^34^, the same stage when full glial coverage and AQP4 polarization are established. Perhaps by not having complete coverage of the brain vasculature by glia^35^, this allows for enhanced plasticity or access to growth factors that facilitate tissue regeneration. Further investigation of glial-vascular interactions in other vertebrates capable of neuronal regeneration, such as salamanders and axolotls, would be necessary to test this hypothesis. Additionally, if zebrafish glia are indeed more ‘immature’ but have the potential to mature into fully ensheathing, mammalian-like astrocytes, it would be intriguing to explore whether promoting this maturation could increase glial-vascular interactions while simultaneously reducing the regenerative capacity of the remarkable zebrafish brain.

## Supporting information

Video S1. Time-lapse imaging of mosaically labelled glia from 3-4 dpf reveals highly dynamic glial projections.

Video S2. Time-lapse imaging of mosaically labelled glia from 6-7 dpf reveals stabilized glial-vascular interactions.

## Acknowledgments

We thank members of the O’Brown and Megason laboratories for data discussions and comments on the manuscript; Dr. Zach O’Brown for discussions and comments on the manuscript; the Zebrafish Core facility at Rutgers for Fish Maintenance. This work was supported by the National Institutes of Health (NIH) R00HD103911 (N.M.O.) and CIHR grant 506513 and U.S. DoD grant 13438153 (B.L.).

## Author contributions

N.M.O., S.G.M. and B.L. conceived the project and designed experiments. N.M.O., C.S., L.G.G., N.P, and M.F.-A. performed experiments. N.M.O and C.S. analyzed the transgenic data and B.L. and M.F.-A. analyzed the EM data. Python image analysis code was developed by B.Z.J. Manuscript was written by N.M.O, L.G.G, C.S., B.L.

## Methods

### Zebrafish strains and maintenance

Zebrafish were maintained at 28.5°C following standard protocols of zebrafish husbandry^19^. All zebrafish work was approved by the Harvard Medical Area Standing Committee on Animals under protocol number 04487 and the Rutgers University Standing Committee on Animal Care under protocol number 202300038. Adult fish were maintained on a standard 12 hour light-dark cycle. Adult fish, age 3 months to 1.5 years, were crossed to produce embryos and larvae. For imaging larvae up until 10 dpf, 0.003% phenylthiourea (PTU) was used daily beginning at 1 dpf to inhibit melanin production. These studies used the AB wild-type strain and the transgenic reporter strain Tg(*kdrl:HRAS-mCherry*)^s896 20^, abbreviated as Tg(*kdrl:mCherry*) in the text.

### Glial Plasmids

Zebrafish glia were labeled by the *gfap:GFP* plasmid^21^, a gift from Pamela Raymond (Addgene plasmid #3971). We generated the *slc1a3b:MYRGFP* plasmid, labeled as *glast:myrGFP* in the text, by excising the P2A-H2A-mCherry cassette, which labeled the nucleus, from the original *glast* plasmid^14^. The *slc1a3b-3kb:MYRGFP* plasmid, labeled as *glastini*:myrGFP in the text, was then generated by concatenating 2 separate parts of the original promoter element using primers 5’-AATATTTTTGTCCGATTGTCTACAC-3’ and 5’-TCTGTCTTCTGCTGCTGTAATCTGT-3’ to amplify the distal peak and primers 5’-AGCAAATGATAATGGTATTAACAGCAGAGAGTTGTATG AAAGC-3’ and 5’-GTGTAGACAATCGGACAAAAATATTAAGGGCGAATTTCGAGCCGGGCC CA-3’ to amplify the rest of the plasmid.

### Mosaic labeling and imaging

Mosaic animals were generated by microinjection of *tol2* mRNA (50 ng/ul) and plasmid DNA (20 ng/ul) at the single-cell stage. Embryos were then raised until the desired stage and fixed with 4% paraformaldehyde in PBS overnight at 4°C. Brains from 1 month old and adult zebrafish were then dissected and cleared with CUBIC as previously described^22^ prior to imaging. Fixed larvae (3-14 dpf) and dissected and cleared brains were mounted in 1% low gelling agarose (Sigma: A9414) on 0.17 mm coverslips and imaged on either a Leica SP8 or Stellaris DMI8 laser scanning confocal microscope at 1024×1024 resolution using the same acquisition settings with a 25x water immersion objective.

### Generation of stable glial transgenic lines and imaging

The Tg(*slc1a3b:MYRGFP*^*nj8*^) and Tg(*slc1a3b-3kb:MYRGFP*^*nj9*^) transgenic lines were generated by raising mosaic animals injected with the respective plasmids and *tol2* mRNA and outcrossing them to AB wildtype animals for 2 generations. Live larvae (3-10 dpf) were immobilized in tricaine and mounted in 1-2% low gelling agarose (Sigma: A9414) on 0.17 mm coverslips and imaged on a Stellaris DMI8 laser scanning confocal microscope at 1024×1024 resolution using the same acquisition settings with a 25x water immersion objective.

### Glial segmentation and analysis

Individual z-stacks were saved as .tiff files and then exported to .HDF5 files using the Fiji Tiff to H5 macro. We then used Ilastik^23^ pixel classification to designate vascular, glial and background signals in 3D. Mosaic animals were then analyzed for glial-vascular contacts and cell morphology using a custom Python script (glial_segmented.ipynb, github:https://github.com/O-Brown-lab/2024GliaContacts). Glial contacts with the vasculature were assessed within a 2 μm dilation of the vasculature. Stable *glast:myrGFP* and *glastini:myrGFP* animals were analyzed using FIJI^24^ by first automatically thresholding the Ilastik segmentations with the Yen method for both the glial and vascular signals. We then used Image Calculator to overlap the thresholded glial and vascular signals and then measured the percent area that was covered in the brain ROI in 200 μm thick maximum intensity projections for both the vascular signal and the overlapping signal.

### Transmission electron microscopy

Fish were anesthetized using tricaine and initially fixed by immersion in a solution of 4% paraformaldehyde (VWR:15713) and 0.1M sodium-cacodylate (VWR:11653). After this quick primary fixation, the adult brains were exposed by removing the dorsal skull. Larvae and adults were then put in a secondary fixation solution (2% glutaraldehyde (Electron Microscopy Sciences: 16320), 4% paraformaldehyde, and 0.1M sodium-cacodylate) at room temperature for 7-14 days. Following complete fixation, the samples were washed overnight in 0.1M sodium-cacodylate. For further preparation, entire larval heads or coronal vibratome sections (50 μm) of adult brains were post-fixed in a solution containing 1% osmium tetroxide and 1.5% potassium ferrocyanide, dehydrated by increasing ethanol concentrations followed by propylene oxide. Afterward, post-fixed sections were embedded in Durcupan ACM Epoxy resin (Electron Microscopy Sciences: 14040). Ultrathin sections (80 nm) were then cut from the block surface, using a Leica EM UC6 ultramicrotome, and collected on copper grids. Samples were examined under JEM-1400Flash transmission electron microscope operating at 80kV and equipped with a 16MP digital camera (GATAN One View). 10 capillaries per animal were randomly imaged. For each capillary, an image at 5600× and 11,000× was acquired. Analyses of the micrographs were performed by a blinded analyzer using ImageJ software. Quantifications of glial endfoot number and coverage were performed as described previously^25^. Individual endfeet were defined as closed areas of dim, electron-clear astroglial cytoplasm in contact with the exterior aspect of the basement membrane (BM) with long, enlarged cisternae and electron-dense mitochondria^26^. The Measure function was used to calculate the area of each endfoot. For each image, the total endfoot number in contact with the vessel was tallied, as was the combined area of all endfeet and the total length of the endothelial BM. All metrics were quantified manually from scaled micrographs in ImageJ (NIH), using tools as Polygon, Freehand Selections or Straight Line, followed by the Measure function.

### Immunogold labeling for electron microscopy

Adult *glast:myrGFP* fish were anesthetized using tricaine and initially fixed by immersion as above. The adults with exposed brains were then placed in a secondary fixation solution with 0.5% glutaraldehyde, 4% paraformaldehyde, and 0.1M sodium-cacodylate for 1 hour at 4°C and immediately washed in cold 0.1M sodium-cacodylate. Coronal vibratome free-floating sections of 50 μm were collected and immersed in 0.1% sodium borohydride/PBS for 30 minutes at room temperature. Samples were thoroughly rinsed in PBS and blocked with 10% goat serum/0.5% gelatin/PBST (0.01% Triton X-100), for 2 hours at room temperature. Sections were then incubated with goat anti-GFP (1:250, Abcam: AB5450) diluted in blocking solution (without Triton X-100) overnight at room temperature. After thoroughly rinsing sections in PBS, sections were then incubated with gold-labeled rabbit anti-goat IgG (1:50, Nanoprobes: 2005-1ML), diluted in blocking solution (without Triton X-100), overnight at room temperature. Sections were washed with PBS followed by 3% sodium acetate rinse (3 × 5 min). Sections were subjected to silver-enhancement, following manufacturer’s instructions (Nanoprobes: 2012-45ML). Sections were then thoroughly rinsed in 3% sodium acetate and extensively washed in 0.1M PB pH 7.4 prior to post-fixation and image acquisition as outlined above.

## Statistical analysis

All statistical analyses were performed using Prism 10 (GraphPad Software). Two group comparisons were analyzed using an unpaired two-tailed t test. Multiple group comparisons were analyzed with one-way ANOVA, followed by a post hoc Tukey’s multiple comparison test. Sample size for all experiments was determined empirically using standards generally employed by the field, and no data was excluded when performing statistical analysis. Standard error of the mean was calculated for all experiments and displayed as errors bars in graphs. Statistical details for specific experiments, including precision measures, statistical tests used, and definitions of significance can be found in the Figure Legends.

## Supplemental Information

**Video S1. Time-lapse imaging of mosaically labelled glia from 3-4 dpf reveals highly dynamic glial projections**.

**Video S2. Time-lapse imaging of mosaically labelled glia from 6-7 dpf reveals stabilized glial-vascular interactions**.

